# Patterns of within-host spread of *Chlamydia trachomatis* between vagina, endocervix and rectum revealed by comparative genomic analysis

**DOI:** 10.1101/2023.01.25.525576

**Authors:** Sandeep J. Joseph, Sankhya Bommana, Noa Ziklo, Mike Kama, Deborah Dean, Timothy D. Read

## Abstract

*Chlamydia trachomatis*, a gram-negative obligate intracellular bacterium, commonly causes sexually transmitted infections (STIs). Little is known about *C. trachomatis* transmission within the host, which is important for understanding disease epidemiology and progression. We used RNA-bait enrichment and whole-genome sequencing to compare rectal, vaginal and endocervical samples collected at the same time from 26 study participants who attended Fijian Ministry of Health and Medical Services clinics and tested positive for *C. trachomatis* at each anatomic site. The 78 *C. trachomatis* genomes from participants were from two major clades of the *C. trachomatis* phylogeny (the “prevalent urogenital and anorecta”l clade and “non-prevalent urogenital and anorectal” clade). For 21 participants, genome sequences were almost identical in each anatomic site. For the other five participants, two distinct *C. trachomatis* strains were present in different sites; in two cases, the vaginal sample was a mixture of strains. The absence of large numbers of fixed SNPs between *C. trachomatis* strains within many of the participants could indicate recent acquisition of infection prior to the clinic visit without sufficient time to accumulate significant variation in the different body sites. This model suggests that many *C. trachomatis* infections may be resolved relatively quickly in the Fijian population, possibly reflecting common prescription or over-the-counter antibiotics usage.

**Importance:** *Chlamydia trachomatis* is a bacterial pathogen that causes millions of sexually transmitted infections (STIs) annually across the globe. Because *C. trachomatis* lives inside human cells, it has historically been hard to study. We know little about how the bacterium spreads between body sites. Here, samples from 26 study participants who had simultaneous infections in their vagina, rectum and endocervix were genetically analyzed using an improved method to extract *C. trachomatis* DNA directly from clinical samples for genome sequencing. By analyzing patterns of mutations in the genomes, we found that 21 participants shared very similar *C. trachomatis* strains in all three anatomic sites, suggesting recent infection and spread. For five participants two *C. trachomatis* strains were evident, indicating multiple infections. This study is significant in that improved enrichment methods for genome sequencing provides robust data to genetically trace patterns of *C. trachomatis* infection and transmission within an individual for epidemiologic and pathogenesis interrogations.

## Introduction

The obligate intracellular bacterium *Chlamydia trachomatis* is the most common worldwide cause of bacterial sexually transmitted infections (STIs) with over 129 million annual cases in 2020(1). In 2019, 1.8 million cases were reported in the United States alone, representing a 19% increase since 2015(2). Approximately 80% of female and 50% of male *C. trachomatis* STIs are asymptomatic(3), increasing the risk of transmission and complications at a yearly cost of billions of dollars(4).

The endocervix is considered the most common initial site of chlamydial sexually transmitted, non-lymphogranuloma venereum (LGV) infections. Sloughed *C. trachomatis* infected cells and the organism itself can be secreted into the vagina but neither infect the squamous epithelium of that organ(3). Cervicitis, an inflammation of the uterine endocervix, is a strong predictor of upper genital tract inflammation and disease(5), including pelvic inflammatory disease, tubal-factor infertility, ectopic pregnancy and poor pregnancy outcomes(6). The rectum is another site of infection. A growing number of studies now show that *C. trachomatis* rectal infections are more common than previously thought, ranging from 2% to 77% of women seen in clinical settings(7). In one study, over 70% of women with urogenital *C. trachomatis* also had rectal *C. trachomatis* infection(8). Of the 24 studies reporting on both urogenital and rectal infections in the same women, six showed a higher prevalence of *C. trachomatis* in the rectum (7). These data suggest that, while the rectum is known to be a common site of infection with LGV strains among men who have sex with men(9), it may also be a more frequent primary site of non-LGV strain infections among women. However, no studies to date have evaluated this issue.

There are several hypotheses for *C. trachomatis* transmission between sexual partners and within anatomic sites of the same individual given our fragmentary knowledge of the genetic structure of *C. trachomatis populations* in natural human infections. The ascertainment of *C. trachomatis* infection in females could be affected by rectal infections persisting longer than endocervical infections and/or increased transmissibility during receptive anal intercourse (RAI). However, a recent study found no association between RAI and rectal *C. trachomatis* infections(10) and another found that screening given a history of RAI did not significantly influence the rate of detection of *C. trachomatis* infections in the rectum(8). We also know that women may develop urinary tract infections from enteric bacteria that are transferred from the perineum or anorectal area during sex(8). It is therefore possible that rectal *C. trachomatis* infections could similarly be spread to the endocervix and urethra. The concern here is that single dose treatment that is effective for uncomplicated urogenital tract infections is inadequate for rectal infections, as has been shown in recent studies(11–15). Indeed, a study that followed cervicovaginal and anorectal *C. trachomatis* loads following treatment with one gram of azithromycin found consistently higher loads in the anorectal site at 16 days after therapy with increasing loads up to 51 days when the study was terminated(16). Due to the requirement to treat non-LGV *C. trachomatis* infections of the rectum for seven days and LGV strains for 21 days, adherence to treatment and/or treatment failure, as a result of lack of adherence, are also concerns(17). These studies show that rectal infection, if not treated appropriately, could have a significant effect on persistence and within-host transmission and disease. Therefore, it is important to understand the pathways of transmission between anatomic sites.

To understand the dynamics and pathobiology of within- and between-host transmission of *C. trachomatis*, we explored the relationships among *C. trachomatis* genomes sequenced using DNA purified directly from endocervical, vaginal and rectal swabs from the same women. Our cohort comprised a population of Fijian women that have an unusually high prevalence of *C. trachomatis* STIs(18). We sought to reveal evidence of within-host dissemination that may promote maintenance of infection in the rectum and increase transmission both within the host and to sexual partners in addition to providing data to select optimal anatomic sites for diagnostic screening, appropriate treatment and duration of therapy.

## Results

### Direct enrichment and sequencing of *C. trachomatis* genomes and comparison of bacterial loads between anatomic sites

Clinical endocervical, rectal, and vaginal swab samples collected from 26 women who attended the Fijian Ministry of Health and Medical Services clinics and tested positive for *C. trachomatis* at each anatomic site simultaneously were supplied de-identified from an ongoing parent study(18) (Supplemental Table 1). We successfully extracted DNA from clinical swabs and used our recently redesigned Agilent RNA bait library(19)to enrich *C. trachomatis* genomic sequences from Illumina sequencing libraries (see Methods). We defined a threshold for a “good quality” genome of at least 10x average *C. trachomatis* genome read redundancy (“coverage”) post-enrichment and at least 5 reads mapped to > 900,000 bases of the 1,042,519 bp *C. trachomatis* reference D/UW-3/CX chromosome. The 26 participants had “good quality” data from all three anatomic sites (78 samples) that were further analyzed in this study (Supplemental Table 1). The median coverage of these 78 samples was 127x with an average of 308x; only three samples were lower than 20x. The RNA bait method was therefore able to enrich *C. trachomatis* genomic DNA even though the samples from the three anatomic sites likely contain high levels of other viral and bacterial organisms. These data are supported by our previous study using the same methodology that successfully generated genomes derived from DNA purified directly from clinical vaginal-rectal pairs from Fijian participants(19).

Using qPCR with conserved *ompA* primers, the chromosomal yield for 25/26 women with *C. trachomatis* successfully sequenced from each body site ranged from 69 to 9,600,000 copies/μL. Given the obligate intracellular nature of *C. trachomatis*, and to normalize against the number of human cells collected in the sample, the ratio of the *C. trachomatis* genomic copy number to the human beta-actin copy number was calculated as an estimated relative load of the organism in each anatomic site. In comparing the vaginal with the rectal site for each woman using a paired t-test, there was a statistically significant higher load in the rectum than the vagina (p = 0.0124; Supplemental Figure 1). However, there were no statistically significant differences between rectum/endocervix and vagina/endocervix sites. When comparing body sites from the same person, 21 of the 26 women had a higher load in the rectum compared to the vagina (Figure 1). However, the differences in qPCR loads across body sites were not reflected in the redundancy of genome coverage. Within the 78 genomes, there was a significantly higher coverage in the endocervical samples compared to rectal (T-test; *P* = 0.031) and vaginal (*P* = 0.0016) samples (Supplemental Figure 2).

**Figure 1.**
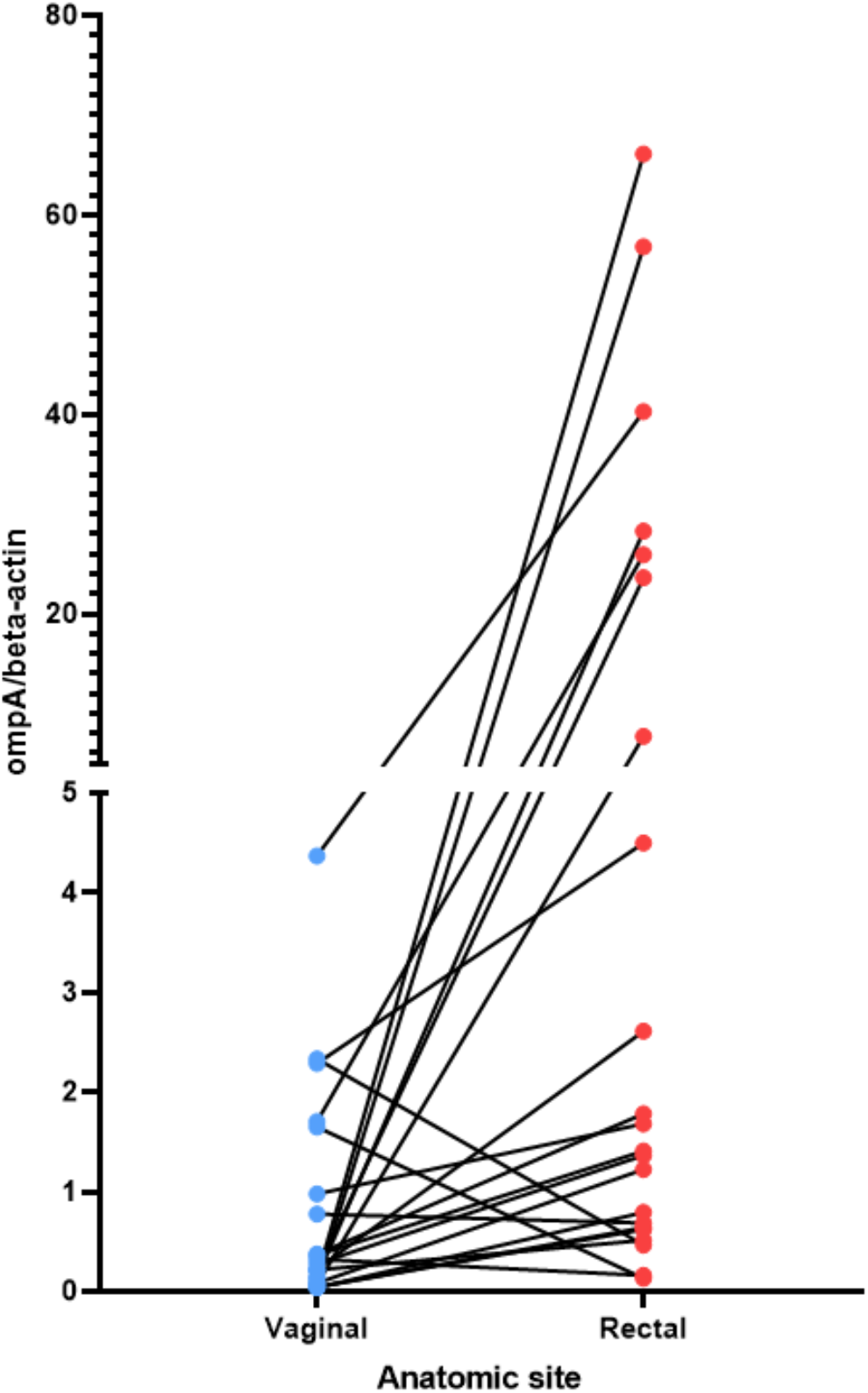
Relative load of *C. trachomatis* in the vagina and rectum estimated by qPCR The non-transformed ratio of the *C. trachomatis omp*A genome copy number to the beta-actin genome copy number is shown (see Methods). C, endocervix; R, rectum; V, vagina. The lines connect the *C. trachomatis* load value for the vagina to the load value for the rectum for the same woman.

### Fiji sample genomes in the context of the global *C. trachomatis* phylogeny

We investigated the phylogenetic distribution of assembled genomes from this study (“Fiji samples”) and selected chlamydial reference and other clinical genomes representing known global *C. trachomatis* clades corresponding to four major *C. trachomatis* clades: LGV, ocular, “prevalent urogenital and anorectal (P-UA) and “non-prevalent urogenital and anorectal” (NP-UA)(20)(Figure 2; Table 1). The reference genome D/UW-3/CX was in the NP-UA clade. All Fiji genomes were in the NP-UA and P-UA clades, forming two subclades of NP-UA and one in P-UA, suggesting that the Fiji genomes were derived from at least two independent introductions in NP-UA and one in P-UA (Figure 2). Based on sequencing of the *omp*A gene, referred to as the *omp*A genotype, 32 genomes in NP-UA had *omp*A genotype D (4), F (3), G (23) and Ja (1) plus one that was not possible to determine, and the 46 genomes in P-UA had E (21), G (2), and Ja (23) *ompA* genotypes. Twenty-four Fiji samples dominated a sub-lineage of NP-UA (*ompA* genotypes G and F) with only one publicly submitted genome sequence: G/11222 (BioSample: SAMN02603694, Assembly NC_017430.1)(21), which was a cervical sample but with no notation of geographic source. This Fijian subclade may represent a local endemic clone. We also found genomes with *ompA* genotype Ja and plasmid genotype E that we had previously described in the Fiji population(19).

**Table 1.** Terms used specific to this work

| Term | Explanation |
| --- | --- |
| CSS | “Clonal SNP sites”. A set of 5,520 SNPs that were used to differentiate Fijian NP-UA from P-UA chromosomal backgrounds. They were defined at positions where 90% NP-UA had one allele and 90% P-UA had the other (reference *C. trachomatis* D/UW-3/CX is in the NP-UA clade). |
| “fixed SNP” | Single Nucleotide Polymorphism is defined here as a position with an allele frequency of less than 0.1 compared to the reference *C. trachomatis* D/UW-3/CX chromosome. For example, if the reference nucleotide at a position is “A”, a fixed SNP would have >90% sequencing reads as either “G”, “C” or “T” aligning to that position. |
| LGV | “Lymphogranuloma venereum”. |
| NP-UA | “non-prevalent urogenital and anorectal” clade in the *C. trachomatis* species phylogeny |
| P-UA | “Prevalent urogenital and anorectal” clade. |
| RAI | “Receptive anal intercourse”. |
| SNV | “Single nucleotide variant”. Defined here as an allele frequency of >0.1-0.9< compared to the reference. |
| STI | "Sexually transmitted infection" |

**Figure 2.**
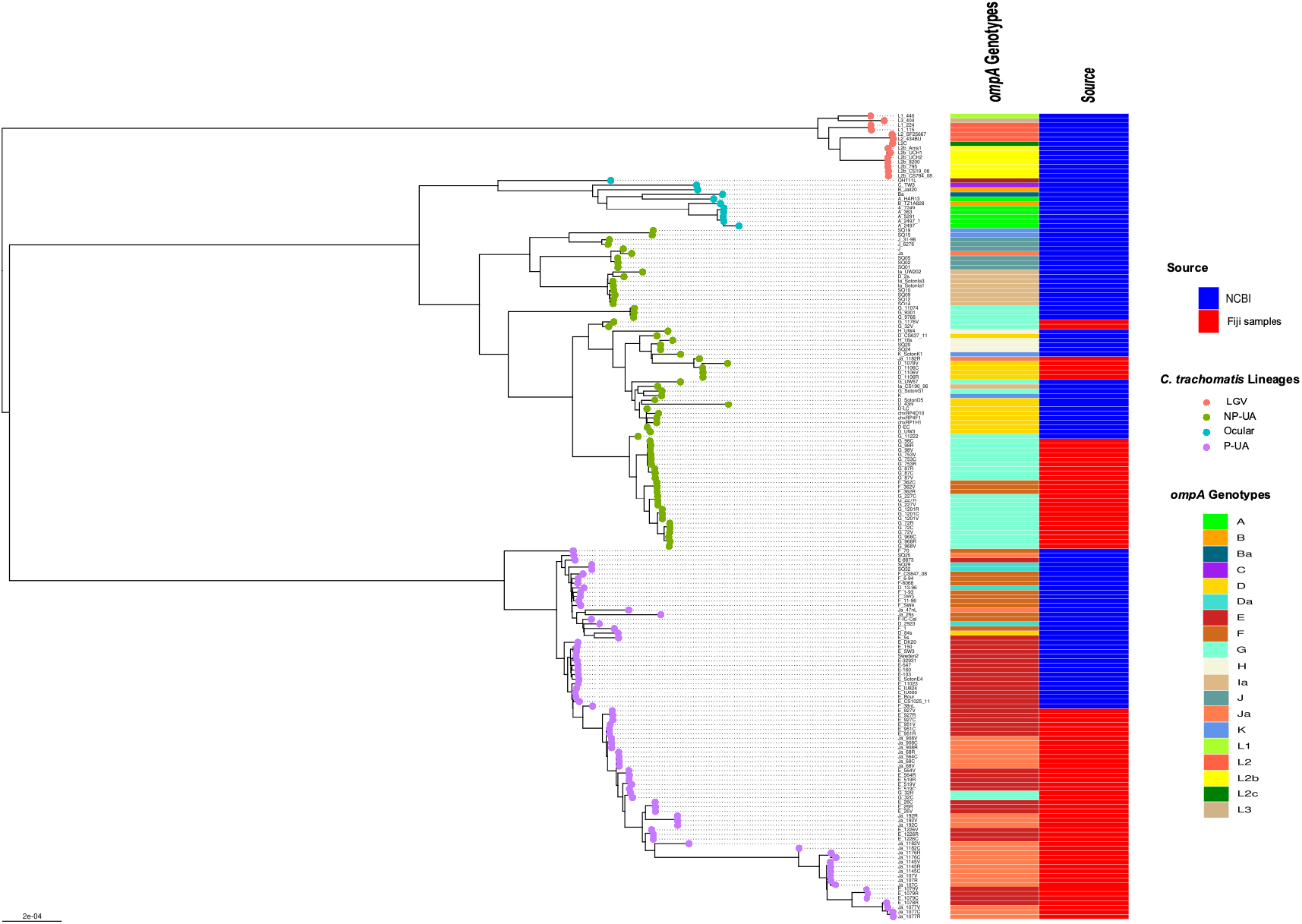
Global Phylogeny with Clade designations The global phylogeny of high-quality *C. trachomatis* Fiji genomes plus selected complete *C. trachomatis* reference and clinical genomes representing global diversity from the National Center for Biotechnology Information (NCBI). Sample names are <*omp*A genotype>-<participant ID>-<body site code, where C = endocervix, R = rectum and V = vagina>. The round tips are colored by the 4 clade designations (LGV, Ocular, Prevalent-Urogenital and Anorectal (P-UA), Non Prevalent Urogenital and Anorectal (NP-UA)). The first column to the right of the tree denotes the ompA genotype with code at the lower right; the second column represents the source of the genomes from NCBI or the Fijian samples.

Numerous studies have shown that *ompA* alleles recombine frequently between *C. trachomatis* genomic backbones(22–26). While the association of *ompA* genotypes with clades in Fiji strains was broadly consistent with patterns found in the Hadfield *et al* study(23), there were some combinations of genomic clade and *ompA* in this work not previously reported: G in P-UA and F in NP-UA (Figure 2). fastGEAR(27) inferred recombination events in ancestors of the global P-UA clade and five (primarily from NP-UA into P-UA) as well as recent recombinational exchange of DNA within the branches of the tree containing Fiji strains (Supplemental Figure 3). Recent inferred events included donors from all clades, including a small number of importation events from LGV and ocular clades, respectively, at recombination hotspots in the chromosome (Supplemental Table 2).

### Participants with samples from three anatomic body sites fell into two groups based on levels of *C. trachomatis* genome diversity

Of the 26 study participants, there was good quality genome sequence data across the three anatomic sites, and 21 had the same *ompA* genotype strain consistent with the rest of its genome that formed a monophyletic clade on the global *C. trachomatis* phylogenetic tree (Figure 2; Supplemental Table 3). We inferred these strains shared a recent common ancestor. We termed these 21 participants “Group A”. Five participants (“Group B”) had three samples that appeared not to derive from a single recent infection event. For participant #1078, the rectal sample and vaginal/endocervical samples were different *ompA* genotypes/genomes from different clades (E in P-UA and D in NP-UA, respectively). For participant #564, all samples were in P-UA but the vaginal and rectal samples were both E while the endocervical sample was Ja and more distantly related on the core genome phylogeny than the other two (Figure 2; Supplemental Figure 3). The rectal and endocervical samples of participant #1176 were both Ja in P-UA, but the vaginal sample was a G in NP-UA. In participant #32, all of the strains were *omp*A genotype G. However, the endocervical and rectal genomes were closely related in P-UA while the vaginal strain was in NP-UA. For participant #1182, all strains were *ompA*genotype Ja but in this case, while the vaginal and endocervical genomes were closely related in P-UA, the rectal genome was in NP-UA. The differences in *C. trachomatis* strains between the vagina and endocervix of the same individual confirm that these sites can be effectively sampled without cross-contamination. In addition, shotgun metagenomics from some of the same samples as in this study also revealed related but diverged communities at each site(28). Further, while the endocervix is the site of infection and secretions along with the infected cells flow into the vagina, the vaginal environment may promote unique pressures on the genomes that are then detected as noted above.

The *C. trachomatis* ~7 kb virulence plasmid was amplified and sequenced in 66/78 samples. For each participant, the genotype based on comparison with reference strain plasmid sequences was identical across the anatomic sites (Supplemental Table 1). All plasmids were in the E genotype or “D/G;” D and G plasmids had identical sequences in our typing scheme. Plasmid genotype E was linked to P-UA genomes (36 out of 37 samples with data) and D/G linked to NP-UA (26/29 samples with data). The strong association between chromosome and plasmid genotype suggested that vertical transmission was the dominant mode for plasmid inheritance(23). Only in Group B patients were incongruent combinations seen (plasmid genotype E-P-UA for 32V, 1176V, and 1182R samples and plasmid D/G-NP-UA for 1078R). These samples likely have had plasmid replacement events, with the donor strain containing the transferred plasmid infecting another anatomic site.

### Patterns of shared fixed SNPs and single nucleotide variants (SNVs) in *C. trachomatis* from anatomic body sites of the same participant are different in Group A and Group B participants

We looked first at the Group A participants to see what the patterns of SNPs revealed about the relationships between the body sites. We defined “fixed” SNPs to mean nucleotide positions on the reference genome where 10% or less of the mapped sequence read coverage matched the reference base. The number of fixed SNPs in all three body sites was 512-1944 for NP-UA samples and for P-UA it was 2169-5229 SNPs (Supplemental Table 3). The higher number for P-UA was because the reference strain D/UW-3/CX was in the NP-UA clade. This pattern was consistent with these SNPs being shared by the common ancestor of the sample that infected the three body sites of each participant.

Fixed SNPs found in only one or two body site samples were rare in Group A participants. The presence of these SNPs would be suggestive of independently evolving populations at different sites. Only five Group A participants had a rectal sample with a fixed SNP, one had a fixed SNP in the endocervix but zero had fixed SNPs unique to the vaginal sample (Figure 3; Supplemental Table 3). One participant had a SNP shared by rectal and vaginal samples and one shared between rectal and endocervical samples. There were three SNPs shared between endocervical and vaginal pairs that were fixed in one of the sites but intermediate frequency in the other (see below)(Figure 3). Since these mutations probably occurred within the host, these data point to a recent common ancestor of the bacterium in each body site of the Group A participants.

**Figure 3.**
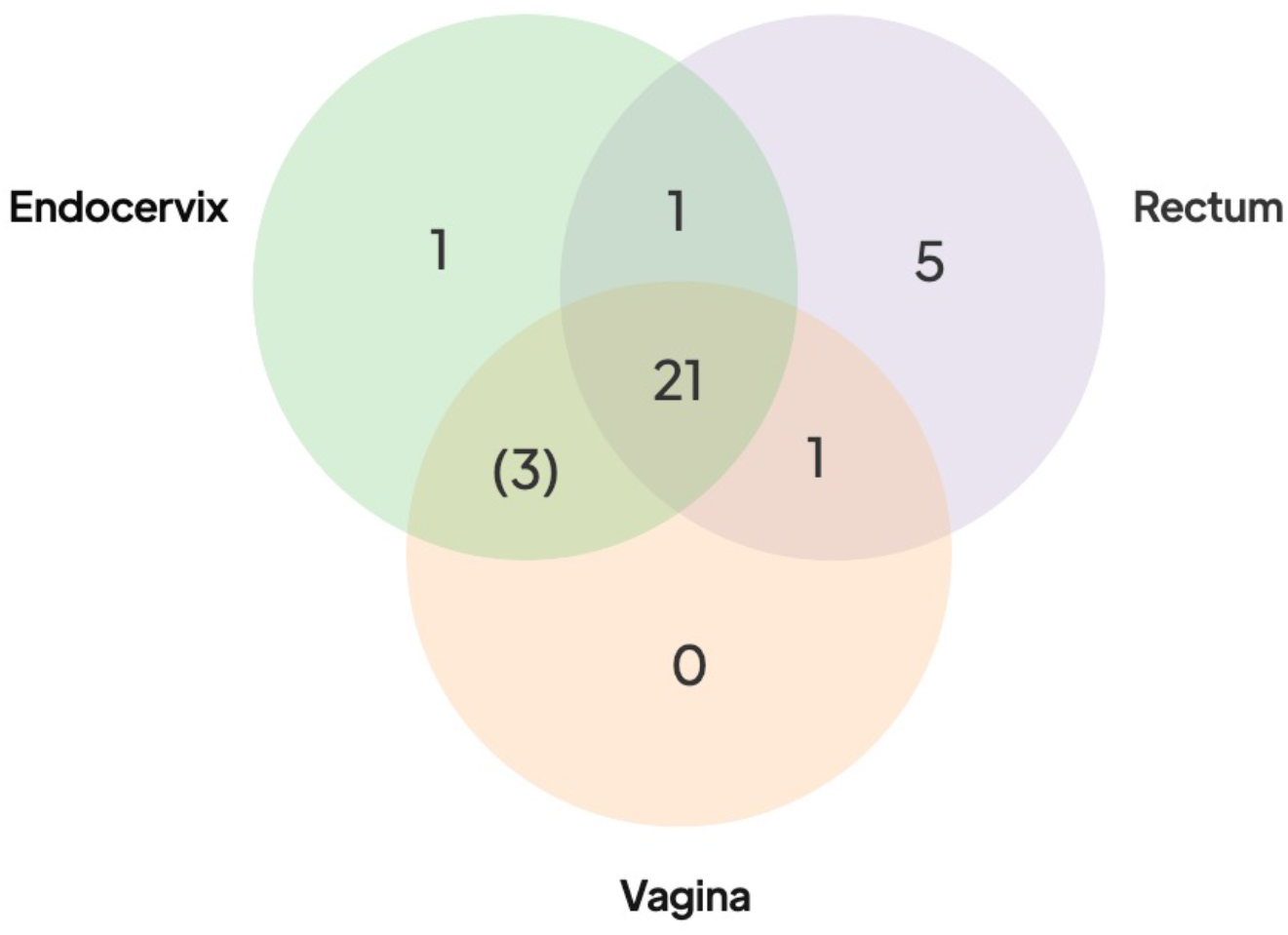
Distribution of shared SNPs by anatomic site in the 21 Group A participants Venn diagram shows the number of participants with fixed SNPs (or fixed in one site with intermediate frequency in the other site in brackets) compared to the reference genome. All 21 participants had shared fixed SNPs in three body sites compared to the reference (center of the Venn diagram). More extensive breakdown of numbers of SNPs by participants are shown in Supplemental table 3.

Next, we looked at intermediate frequency single nucleotide variants (SNVs), which we defined as having reference allele frequencies in the 10-90% range. Reference alleles over 90% were inferred to be the same as the reference while less than 10% were defined as fixed, as in the paragraph above. Across the 78 high-quality Fiji genomes, we found 8,694 SNVs. Most SNVs were found in a small number of samples, with 2,039 (23.5%) found in only one. Of particular note were the 3,818 “rare SNVs” that were only found in one or more anatomic sites of the same participant. The remaining “common” SNVs (found in 4+ of the 78 samples) appeared to be frequently occurring polymorphic sites within *C. trachomatis* populations. They were distributed across the genome but there was a peak in regions around the highly recombinogenic *ompA* gene. SNVs could be generated by genetic drift and/or sharing of populations between body sites. Alternatively, they could be artifacts of random sequencing error. Artifacts would be more likely to occur where there was lower coverage, as one or two miscalled bases could put the position in the 10-90% range for SNV calling. Some Group A participants with lower coverage had as many as 500+ SNVs in only one of the body site samples but on inspection we found that SNVs at these positions were close to the 90% reference threshold, suggesting that they were likely to be false positives generated by sequencing error. Positions that had SNPs that were either fixed in two sites and SNV in the other, or fixed in one and SNV in the other two also were likely artifacts. In this case the SNVs were found at the 10% threshold and probably represented false positive SNPs that were fixed in all three sites. However, positions that were fixed SNPs in one body site but SNV at one other would be expected to be generated infrequently by sequence error. This pattern only occurred in three participants where, in each case, the body site that shared the mutations were the endocervix and vagina (Figure 3).

To help understand patterns of sharing within individuals we identified 5,520 genome positions that differentiated NP-UA and P-UA Fiji strains (see Methods). Because of pervasive recombination in *C. trachomatis* every strain had some alleles assigned to both clades but were overrepresented in alleles common in their own clade. In Group A samples, these clonal SNP sites (CSS) segregated across the chromosome as fixed differences (i.e., either mostly >90% or <10% reference allele frequency). The pattern seen in participant #1201 (Figure 4) is representative of the simple relationships seen in Group A. In this case, CSSs were dominated by NP-UA alleles (> 90% reads aligning to reference bases) with few intermediate frequency SNVs. In Group A participants where the dominant strain was from the P-NP clade, the majority of CSS alleles were different from the reference genome (<10% reads aligning).

**Figure 4.**
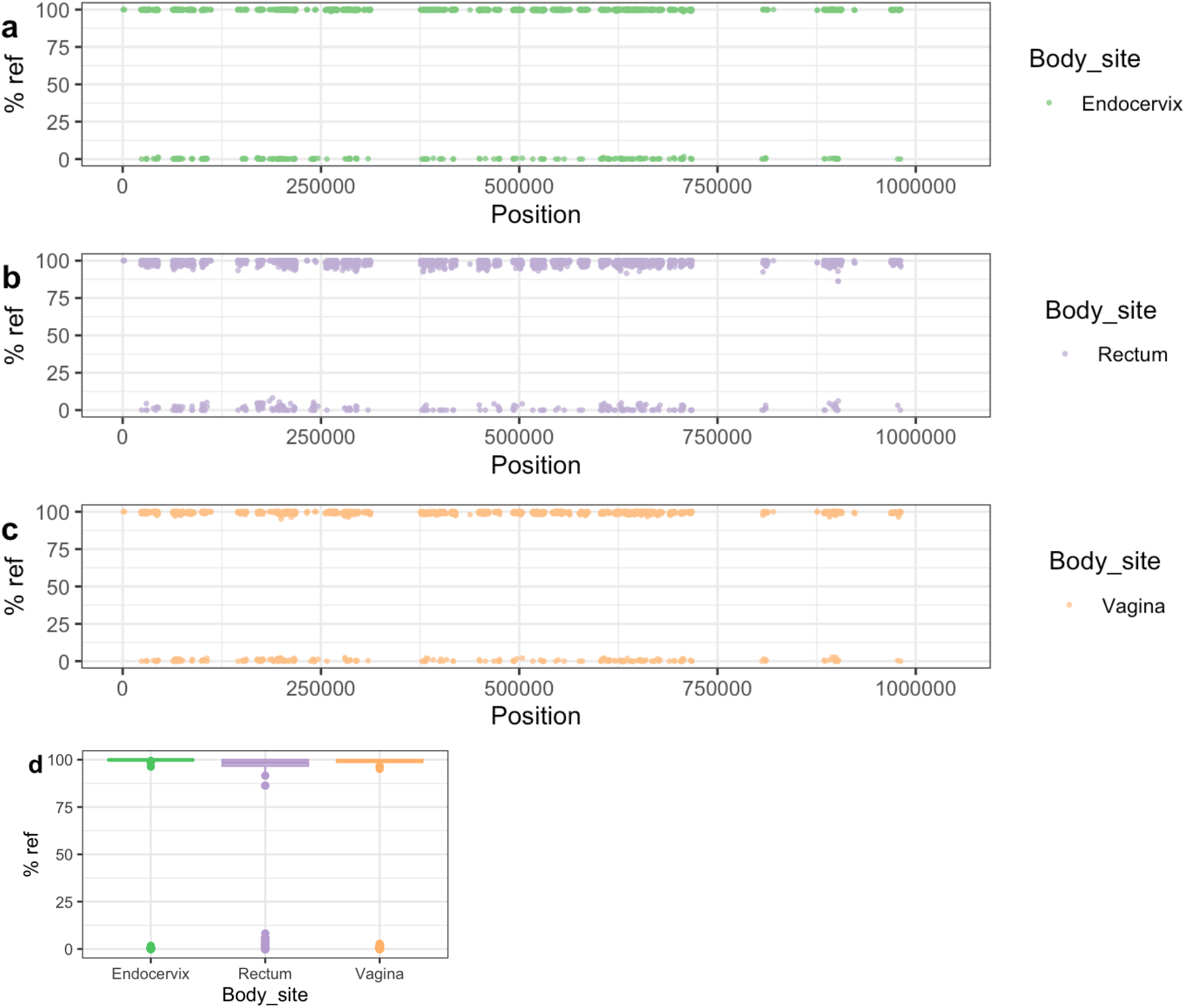
Patterns of SNP and SNV frequency across anatomic sites for representative Group A participant #1201. (a-c) Percent reference scores versus position on reference genome for “clonal SNPs” (CSSs) by body site. The set of 5,520 CSSs were chosen to differentiate NP-UA and P-UA genetic backgrounds. Each point shows the percentage of reads that mapped with the reference allele at each CSS position. The strains from #1201 are from the NP-UA clade and therefore, at most, CSSs are close to 100% match to the reference D allele, which is also in the NP-UA clade. The gaps in the distribution of CSSs across the chromosome are where there were regions of low variation or high recombination. (d) Box plot of distribution of % reference for clonal SNPs by body site. The minority of the CSSs with alternative alleles (<10% of reference genome) were likely the product of recombination events that have occurred since the divergence of the strains. Notably there is an intermediate frequency of SNVs.

We saw more complex patterns of SNPs and SNVs in Group B participants compared to Group A. The simplest Group B participant was #564 where all genomes were in P-UA: the rectal and vaginal genomes were genotype E while the endocervix was genotype Ja. Therefore, the CSS showed all three sites exhibited a pattern typical of P-UA but the rectal and vaginal samples shared a large number of fixed SNPs (179 SNPs) not found in the endocervical sample. Conversely, the endocervical sample had unique fixed SNPs (231 SNPs) not found in the other two body sites (Supplemental Table 3; Supplemental Figure 4). Approximately 50% of these unique SNPs were found within blocks predicted by fastGEAR, suggesting that recombination was a major contributor to genetic differences between the two strains. A simple explanation of these patterns was that participant #564 contained multiple strains: caused by a P-UA Ja strain coinfecting the endocervix after another P-UAE strain had previously infected the rectum and vagina; the reverse order, with E strains coinfecting was also possible. The recombination events between genotypes could have occurred pre- or post-coinfection as natural transformation only requires that chlamydial DNA from a prior, non-viable infection or co-occurring infection be present that can be taken up by a newly infected cell.

In participants #32 and #1176, CSS patterns clearly showed strain mixing in the vaginal genome (Figures 5 and 6). While the endocervical and rectal genomes were dominated by alleles typical of P-UA strains, the vaginal genomes, located in the NP-UAs clade, had intermediate allele frequency across the length of the chromosome. Our interpretation of this pattern is that the vaginal samples contain a mixture of strains with P-UA and NP-UA chromosomes.

**Figure 5.**
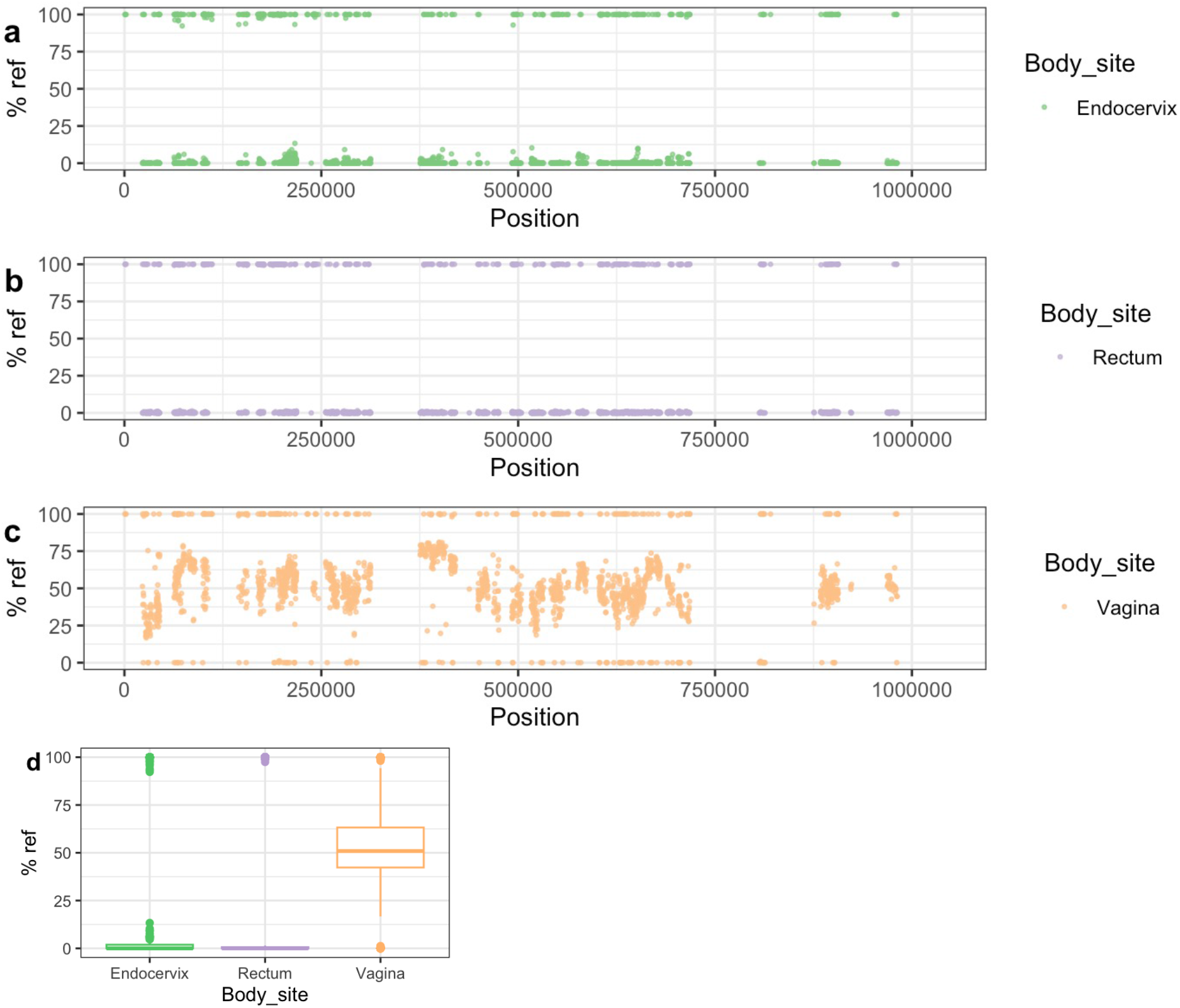
Patterns of SNP and SNV frequency across anatomic sites for Group B participant #32. See legend for Figure 4. The endocervical and rectal strains were in the P-UA clade and therefore the majority of the CSSs had an alternative allele (<10% reference genome). The vaginal genome showed intermediate allele frequency across the chromosome, which was evidence of mixture between P-UA and NP-UA strains

**Figure 6.**
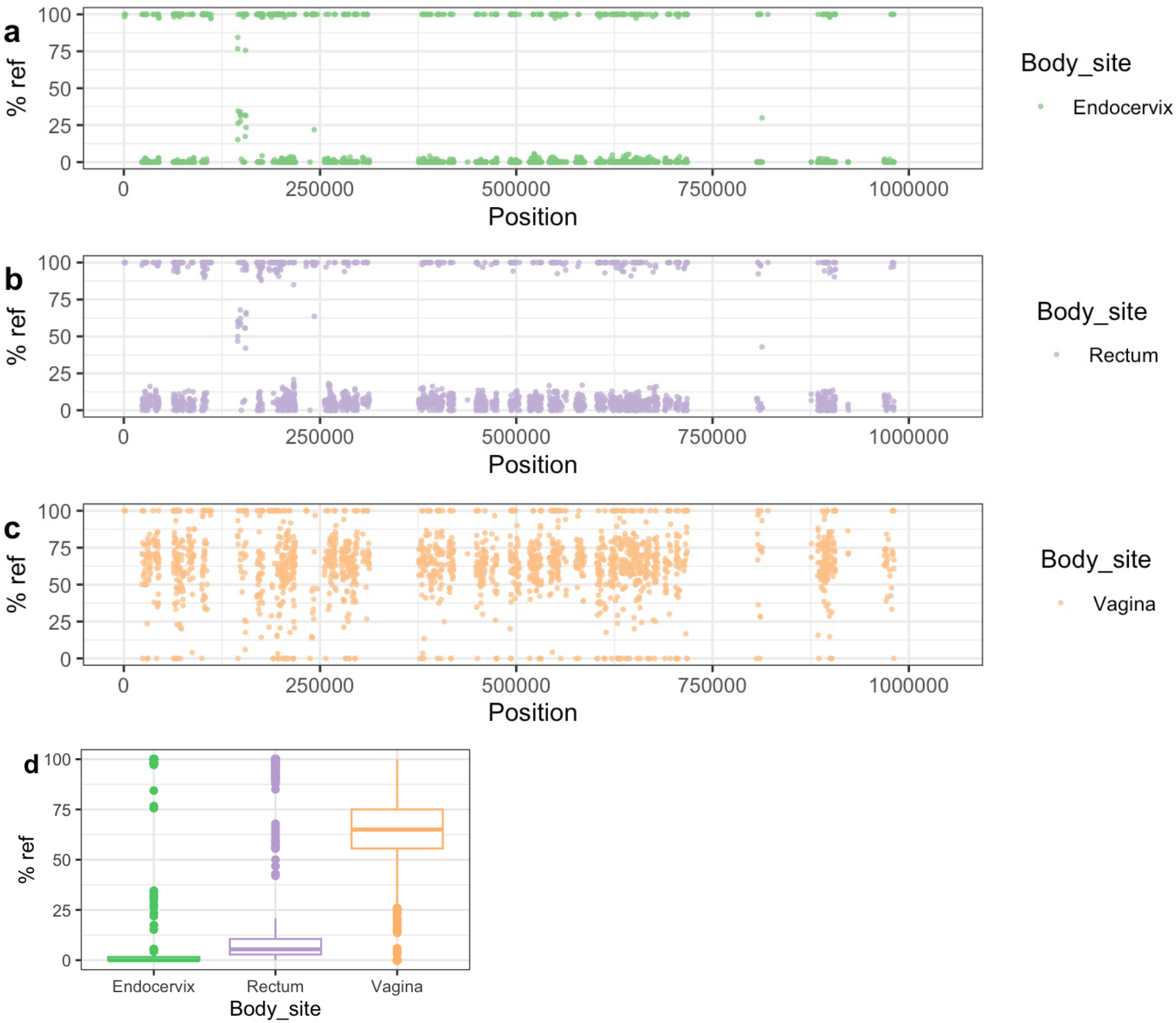
Patterns of SNP and SNV frequency across anatomic sites for Group B participant #1176. See legend for Figure 4. The patterns in this participant are similar to Figure 5 in showing evidence for the vaginal strain being a mixture of P-UA and NP-UA strains.

In participant #1078 interpretation was complicated by the lower data quality of endocervical and rectal samples: (only 249,618 and 875,018 bases with > 10x redundancy, respectively) (Supplemental Figure 5). There were many fixed SNPs in the rectal E genome, indicating it was from a different clade to the vaginal D genome. The pattern of CSS suggested some mixing of P-UA and NP-UA backgrounds in the vaginal genome. In #1078 all samples had the same plasmid subtype “D” despite differences in chromosome backgrounds suggesting possible plasmid transmission.

## Discussion

The Fijian genomes sequenced in this study represent a sampling from the globally distributed P-UA and NP-UA clades. Although the phylogeny suggested multiple introductions of *C. trachomatis* strains from outside Fiji, there was also evidence for clonal expansion in both clades, presumably due to endemic local transmission(18). There was evidence of recent DNA exchange between P-UA and NP-UA clades and possible local introductions of DNA from LGV and ocular clades into both clades of Fijian strains. This suggests that LGV and ocular strains might be present locally in Fiji but not common in the cohort we sampled, which would be expected as LGV is more common in Men who have Sex with Men (MSM), and the ocular strains are associated with the non-STI disease trachoma. Trachoma is endemic to the Pacific Islands of the Western Pacific Region, which would provide an opportunity for exchange during eye infections with both ocular and urogenital strains(29).

This work is centered around sequencing *C. trachomatis* genomes directly from clinical samples using Agilent RNA bait libraries. This approach has been used to sequence bacterial species such as *C. trachomatis* and *Treponema pallidum* that are difficult to culture and are present in only small fractions of the metagenome(19, 23, 24, 30–32). Here, we showed that the newly redesigned bait library(19) could be used efficiently to produce high-quality genome sequences from samples with low yields of *C. trachomatis*, as measured by qPCR. Some samples had a high proportion of human DNA even after enrichment, leaving lower-coverage regions in the *C. trachomatis* genomes. However, we achieved good sequence data from 78 samples representing all three anatomic sites from 26 study participants.

There were complexities in the bioinformatic interpretation of the data, which arose because what was being sequenced was actually a within-species pool of strains rather than the pure cultures normally used in bacterial genomics projects. We showed that SNVs, which we defined here as having an allele frequency of 10-90%, were common across all samples but unevenly distributed, with some having many thousands more than the average due to random sequence errors in low-coverage regions. Non-artifactual SNVs could theoretically come from two sources: 1) the presence of more than one *C. trachomatis* strain through mixed infections; 2) mutation accumulation over time through population growth. Interpretation of the sequence pools in the absence of being able to culture pure cell lines is complex as multiple processes may be occurring, especially allowing for the possibility of recombination between subpopulations of strains within the sample pool(26). We also found that SNVs complicated analysis based on calling consensus nucleotide positions (e.g., *de novo assembly* or reference mapping using tools such as SNIPPY). While these methods worked well for placing samples on a phylogenetic tree, detailed analysis can be confused if SNVs are around the 50% consensus line. Consensus base calling means that distinct subpopulations are not recognized if they are distributed at significantly less than 50% frequency or alternatively, if over 50%, they are incorporated into the consensus.

To our knowledge, this is the first study to use genomics to assess within-host transmission dynamics for *C. trachomatis* STIs. Our analysis revealed two strikingly different patterns within participants: “Group A” (n=21) had three anatomic samples with similar genetic background and *ompA* genotype, while “Group B” (n=5) had one sample with a different background, implying a coinfection event. In the case of Group A, it was notable that only a minority of participants had samples with any fixed SNP differences and, if present, the modal number of SNPs was one (Figure 3; Supplemental Table 3). We argue that positions that were SNPs or rare SNVs shared between two samples from different anatomic sites were likely to be real. These, too, were rare in Group A women (Figure 3). As the mutation rate of *C. trachomatis* inferred from dated whole genome comparison is ~0.2 SNPs per genome per year(23, 24), the most likely implication of these patterns is that there has been recent acquisition and transmission between anatomic sites in these participants. The simplest explanation is that these infections are quite transient and resolve before there has been time to accumulate significant variation between sites. This resolution may be due to recent infection and prescribed treatment proximal to a clinic visit or self-treatment with antibiotics that are available over-the-counter, limiting the longitudinal acquisition of SNPs. Either of these scenarios could result from symptomatic infection and health care seeking behavior or asymptomatic infection with concern over sexual exposure to someone with an STI. These patterns could also be explained by more complex alternative models, for example, population contractions across all body sites followed by rapid re-seeding from one site with a small bottleneck.

The patterns of mutation might reveal pathways of transfer of *C. trachomatis* between anatomic sites, although care must be taken to not over-interpret the findings as the number of participants in this pilot study was small. SNPs and SNVs have been used to infer transmission between individuals(33), and in theory could also be used for potential events occurring between body sites of the same individual. It is possibly a sign of the biases in transmission between sites that unique fixed SNPs in Group A participants were more common in rectal samples, and that vaginal and endocervical samples more often had shared fixed and SNVs (Figure 3). The accumulation of SNPs in one site could be seen as a sign of population stratification caused by anatomy: The vaginal and endocervical *C. trachomatis* populations transmit between each other more frequently, given their proximity, than *C. trachomatis* in the rectum.

The Group B participant samples had much greater numbers of fixed and intermediate SNPs in pairwise comparisons than Group A. The simplest explanation for these is the coinfection of one anatomic site. The site with the divergent strain was not constant: In two cases it was the rectum (participants #1078R and #1082R), in two cases the vagina (participants #32V and #1176V) and in one case the endocervix (#564C). In four of these samples (#32V, #1078R, #1176V, #1182R) there was evidence of mixtures between *C. trachomatis* strains from different clades. These data show that an sequencing of enriched *C. trachomatis genomes* directly from DNA of clinical samples can be used to identify co-infections, which are necessary for inter-strain recombination events to occur. The harmonization of plasmid genotypes in women containing *C. trachomatis* from different clades suggested that the process of plasmid replacement can be rapid. However, the caveat is that plasmid sequences in this study are based on PCR amplification and Sanger sequencing rather than Agilent bait pulldown, so it may not be possible to identify minor plasmid subpopulations.

This study revealed the intricacies of *C. trachomatis* within-host diversity and transmission during natural human infections and suggested that further investigation will yield information that will help understand infection spread and disease processes. More samples are needed from a global sample set to know if these results can be extrapolated across human populations. Integration with bio-behavioral data will also be important to fully understand causes and direction of *C. trachomatis* transmission. Although it would be ideal to expand individual datasets by conducting longitudinal studies to help resolve the dynamics of recombination and determine if multiple cycles of cross-infection occur between sites, this would not be ethical as identification of infection requires treatment to eradicate *C. trachomatis*. Genomic approaches that resolve the potential subpopulations, such as single-cell sequencing(34) and Hi-C(35), are hampered by *C. trachomatis* being only a minor component of the DNA in the clinical metagenomic sample. It may be possible to dissect recombination by isolating clonal *C. trachomatis* populations from individual samples and sequencing them independently. The technique commonly used for this is the plaque assay that is labor-intensive and not always guaranteed to completely separate out subpopulations(36). The most productive near-term strategy may be to continue to build up our picture of *C. trachomatis* natural infection by taking more “snapshots” of populations at single time points across multiple anatomic sites from a larger sample sizes of participants across Fiji, using the efficient RNA-bait methodology, to see if the patterns hold or diverge across a more global population, especially as tourism is a major part of the economy in Fiji.

## Methods

### Study design and Sample Collection

The parent study was cross-sectional in design, enrolling women 18 years of age and older attending Ministry of Health and Medical Services (MoHMS) Health Centers in Fiji following written informed consent as described(18). Appropriate IRB approval had been obtained from UCSF (21-33864) and the Fijian MoHMS (FNHRERC 2015.100.MC) prior to commencement of the parent study. The current study was supplied with *C. trachomatis* positive endocervical, vaginal and rectal swab samples that had been de-identified with a unique ID number. All endocervical samples were collected by trained clinicians after cleaning the exocervix with a large cotton swab prior to inserting the collection swab directly into the endocervix, avoiding contact with the exocervix, vaginal wall or speculum. In addition, data on age were provided at the time of sample collection, and none of the women reported anal intercourse.

Paired vaginal and rectal swabs were screened for *C. trachomatis* using the Cepheid Xpert CT/NG assay (Sunnyvale, CA) according to manufacturer’s instructions. *C. trachomatis* positive endocervical samples were identified using a *C. trachomatis*-specific in-house qPCR assay as described(19).

### DNA extraction and determination of *C. trachomatis* copy number and load

Genomic (g)DNA was extracted from remnant Xpert CT/NG transport media for vaginal swabs and remnant M4 transport media (Thermo Fisher, South San Francisco, CA) for endocervical and rectal swabs as described previously(19). Briefly, 59 μl consisting of 50 μL lysozyme (10 mg/mL; MilliporeSigma, St. Louis, MO), 3 μl of lysostaphin (4,000 U/mL in sodium acetate; MilliporeSigma) and 6 μl of mutanolysin (25,000 U/mL; MilliporeSigma) was added to 200 μl of remnant transport media and incubated for 1 hour at 37°C as described (59). The QIAamp DNA mini kit (Qiagen, California) was then used for DNA extraction, according to manufacturer’s instructions. 5μL of the resulting DNA underwent one or more displacement amplifications using the Repli-G MDA kit (Qiagen), to enrich microbial DNA. DNA concentration was measured using the Qubit dsDNA broad-range assay kit (Invitrogen).

Quantitative PCR (qPCR) was used to determine *C. trachomatis* genomic copy number and *C. trachomatis* load as described(37, 38). Primers specific for the *C. trachomatis ompA* gene and for human Beta-Actin were used to generate standard curves of 10-fold serial increases in plasmids containing a single copy of each gene, respectively. Copy number of *C. trachomatis* and Beta-Actin for the clinical sample was determined based on comparison with the standard curve for the respective control plasmid. *C. trachomatis* load was estimated based on the ratio of bacteria (*C. trachomatis* genome copy number) per human cell (Beta-actin genome copy number) for each clinical sample to normalize the data against the host cell.

### *C. trachomatis ompA* genotyping and plasmid sequencing

The *omp*A genotype was determined for each clinical sample as described previously(36). PCR was performed using primer pairs that flank the *omp*A gene; the product was sequenced in both directions and aligned using MAFFT v7.45062 to create the consensus sequence, which was then aligned with the 19 known *C. trachomatis* reference sequences to determine the *omp*A genotype. The reference strains were A/HAR-13, B/TW-5/OT, Ba/Apache-2, C/TW-3/OT, D/UW-3/Cx, Da/TW-448, E/Bour, F/IC-Cal-13, G/UW-57/Cx, H/UW-4/Cx, I/UW-12/Ur, Ia/UW-202, J/UW-36/Cx, Ja/UW-92, K/UW-31/Cx, L1/440, L2/434, L2a/UW-396, L2b/UCH-1/proctitis, L2c, and L3/404.

The plasmid for each clinical sample was sequenced as described(19). Five primer pairs that flanked and covered the entire plasmid sequence were used, and the PCR products were sanger sequenced and aligned as above using MAFFT v7.45062(39). Each plasmid sequence was aligned to the 19 reference sequences to determine the plasmid identity.

### Enrichment of *C. trachomatis* sequences from clinical samples using an Agilent bait library

We used a methodology for RNA bait capture of *C. trachomatis* described in detail by Bowden et al(19). Human gDNA (Promega, San Luis Obispo, CA) was added to the extracted gDNA from the clinical swabs to reach a total input of 3 μg/130uL for fragmentation and library prep. Samples were sheared on the Covaris LE220 plus (Covaris, Woburn, MA). After shearing and magnetic bead purification, the SureSelectXT Target Enrichment System for Illumina Paired-End Multiplexed Sequencing Library (VC2 Dec 2018) and all recommended quality control steps were performed on all gDNA samples. The 2.698 Mbp RNA bait library consisted of 34,795 120-mer probes spanning 85 GenBank *C. trachomatis* reference genomes(19)(Agilent Technologies, INC, Santa Clara, CA, reference: ELID: 3173001). A 16-hour incubation at 65°C was performed for RNA bait library hybridization. Post-capture PCR cycling was set at 12 cycles based on a capture library size > 1.5 Mb. The libraries were paired end sequenced for 150 nt using an Illumina HiSeq instrument. Sequence data from this project was submitted to the NCBI Sequence Read Archive under the BioProject accession ID: PRJNA609714

### Post-sequencing bioinformatic isolation of *C. trachomatis* sequences

The post-enrichment raw sequencing reads were processed to remove the host genome and *C. trachomatis reads* were extracted and assembled into contigs as described in(19). We used an arbitrary threshold for good quality sequence data if the samples had at least 10x average *C. trachomatis* genome coverage post-enrichment and at least 5 reads mapped to > 900,000 bases of the 1,042,519 Mbp *C. trachomatis* reference D/UW-3/CX chromosome. To genotype the patient samples, de novo contigs were used to extract and compare the *ompA* genes against a customized BLAST(40) database of the 21 reference *ompA* sequences as we described(19).

### Phylogeny and recombination inference

For the global phylogenetic analysis of the main chromosomes (total n= 176), we included all “good quality” genome sequences from the 26 participants (n=77, with the exception of 1078C, which assembled into too many small contigs); and a collection of diverse C. *trachomatis* chromosomes available in NCBI (n=99). We used a reference mapping approach with a custom version of *C. trachomatis* D/UW-3/CX by masking the 6 rRNA genes present in the repeated rRNA operons as described in(19), and generated a full-length whole genome alignment using snippy v4.3.8 ((https://github.com/tseemann/snippy). Snippy mapped the *C. trachomatis* reads from each sample to the reference genome using bwa and identified variants using Freebayes v1.0.2(41). The length of the region common to all samples with at least 10X read coverage and 90% read concordance at each site was 699,239 nucleotides with 11,971 polymorphic sites. Regions of increased density of homoplasious SNPs introduced by possible recombination events were predicted iteratively and masked using Gubbins(42). The final maximum-likelihood (ML) global phylogenetic tree on 10,045 polymorphic sites was reconstructed using RAxML v8.2.9(43) on the recombination removed (MRE) convergence criterion, along with ascertainment bias corrected using Stamatakis method. Lineage-specific phylogenetic trees were inferred as described above by using only the genomes from Fiji samples from their respective lineages.

fastGEAR(27) was run on a whole alignment that contained all “good quality” Fiji *C. trachomatis* genomes along with representative reference genomes from the clade on the global phylogenetic tree. This software infers the population structure and detects the “ancestral” and “recent” recombinations between the genomes present in the alignment. FastGEAR was run by clades with 100 iterations and checking for convergence. The statistical significance of the inferred recombination events (changes in SNP density between the two lineages) were assessed based on the natural log of Bayes factor calculated within FastGEAR. To understand the recombination events within group A individuals, we generated individual whole genome alignments from each of the three body sites by reference mapping the *C. trachomatis* reads to *C. trachomatis* D/UW-3/CX genomes using snippy and the within individual recombination events were inferred using Gubbins as described above.

### Comparison of SNPs patterns between samples from the same participant

We used samtools mpileup(44) to process the BAM files created by aligning sample FASTQ files against the reference chromosome to create tables of the numbers of each base (A, C, T, G) mapped to each individual base of reference. For each pair of samples from the same participant, we used R tidyverse tools(45, 46) to merge the positions with at least 10x read mapping redundancy. Code for analysis of the merged mpileup output was deposited to GitHub (https://github.com/Read-Lab-Confederation/Ct_MAP_analysis).

To create a list of clonal SNP positions (CSSs), we performed Snippy alignment of all contigs from Fiji samples against the reference and identified positions where at least 90% of P-UA strains were identical but different to at least 90% of NP-UA strains. We then filtered out those falling in recombinant regions identified by Gubbins (see section above), leaving 5,520 CSS positions.

## Acknowledgements

We thank the parent study for providing the de-identified samples and for this study and Fijian colleagues: Rachel Devi, Kinisimere Nadredre, Mere Kurulo, and Darshika Balak. Thanks to Brian Raphael and Ellen Kersh for reading through the manuscript. TDR and DD were supported by United States National Institutes of Health award AI138079. The findings and conclusions in this report are those of the authors and do not necessarily represent the official position of the Centers for Disease Control and Prevention. We declare no competing interests.

## Supplemental Material

**Supplemental Table 1**. Metadata, typing and genome sequence quality control statistics associated with the samples from 26 women with good quality genomic sequences in three anatomic sites.

KEY: Age, participant age when samples were taken. Symptoms, whether or not the anatomic site was showing any signs and/or participant had symptoms suggestive of a sexually transmitted infection.

**Supplemental Table 2**. Coordinates of recent cross-clade recombination events inferred by fastGEAR.

KEY: Start and End are the coordinates of the putative recombination region on the reference chromosome. The donor clade codes are as described in Figure 2. In some cases, the donor is unknown or uncertain, probably representing unsampled lineages of *C. trachomatis*. logBF is the log of the Bayes Factor score. Fiji genome names are from Supplemental Table 1. *omp*A genotypes are provided for each Fiji genome.

**Supplemental Table 3**. Sample information from Group A and B participants. KEY: “Fixed SNPs are defined as < 10% reference allele frequency and not in fastGEAR defined recombination blocks. SNVs have 10-90% reference allele frequency. “Rare” SNPs or SNVs are only found in <= 3 samples, which in almost all cases means that they only appeared in samples isolated from one study participant.

**Supplemental Figure 1**

Distribution of log-transformed ratio of the *C. trachomatis ompA* genome copy number to the beta-actin genome copy number (y-axis) is shown for each site. The load was significantly higher in the rectum compared to the vagina (*P* = 0.0124). C, endocervix; R, rectum; V, vagina.

**Supplemental Figure 2**

Comparison of the mean depth of sequencing coverage based on mapping of quality trimmed reads to the reference genome. Significant differences for endocervical depth compared to rectal and vaginal depth are shown. C, endocervix; R, rectum; V, vagina.

**Supplemental Figure 3**

Whole genome phylogenies of strains from this study from clades a) P-UA and b) NP-UA. Two G strains were found in the P-UA clade while three F and four D strains were found in the NP-UA clade. P-UA, prevalent urogenital and anorectal; NP-UA, non prevalent urogenital and anorectal

**Supplemental Figure 4**

Patterns of CSS frequency across anatomic sites for participant 1078. For details of the plots see Figure 4. In this case, strains from the endocervix and vagina were in the NP-UA clade, and the rectum in the P-NP clade.

**Supplemental Figure 5**

Patterns of CSS frequency across anatomic sites for participant 564. For details of the plots see Figure 3. The high number of SNVs seen in this strain were a result of random errors in low sequence coverage regions.

