## Supplementary figures and images for "Patterns of within-host spread of *Chlamydia trachomatis* between vagina, endocervix and rectum revealed by comparative genomic analysis"

### Supplemental Figure 1

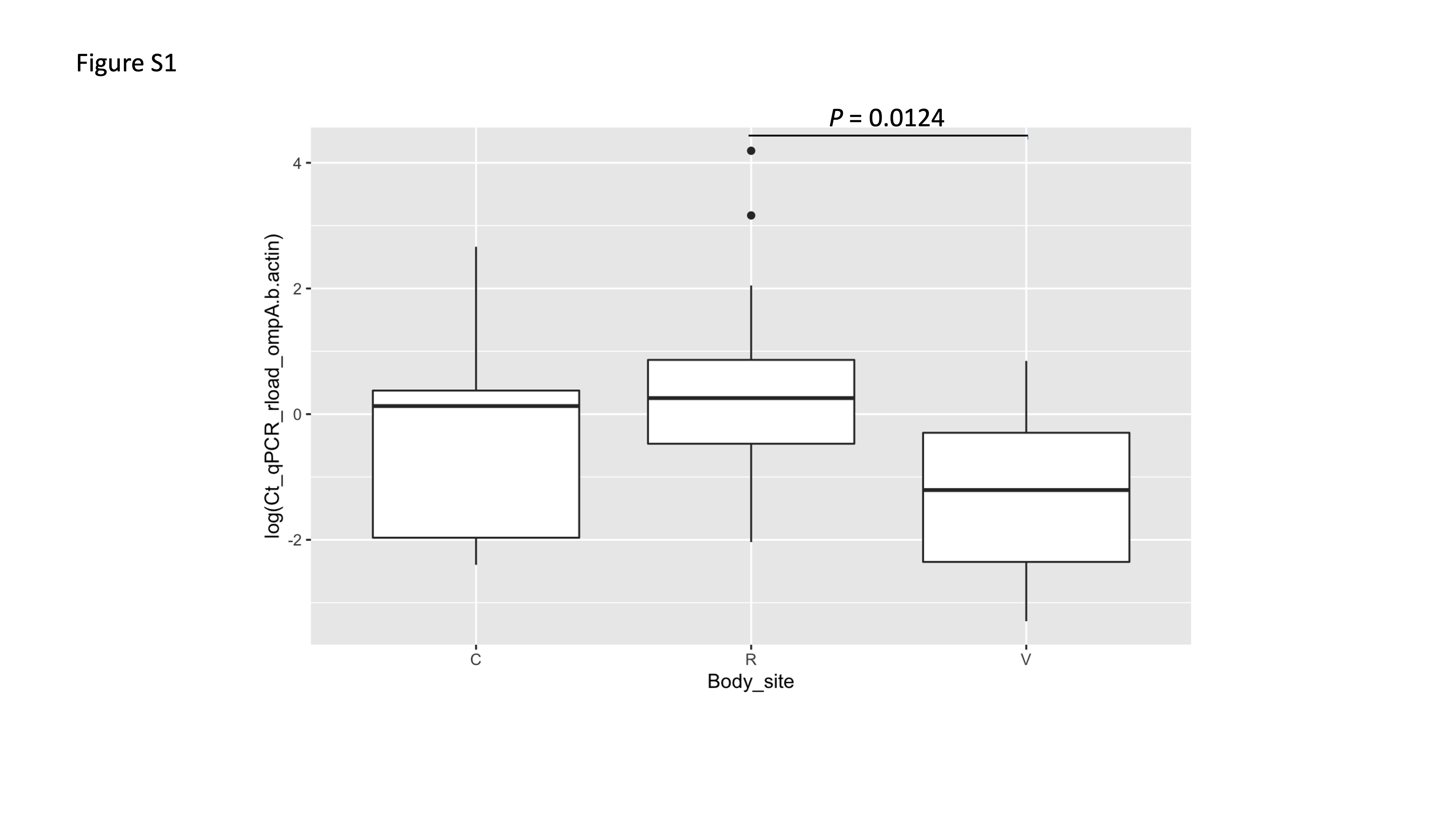

### Supplemental Figure 2

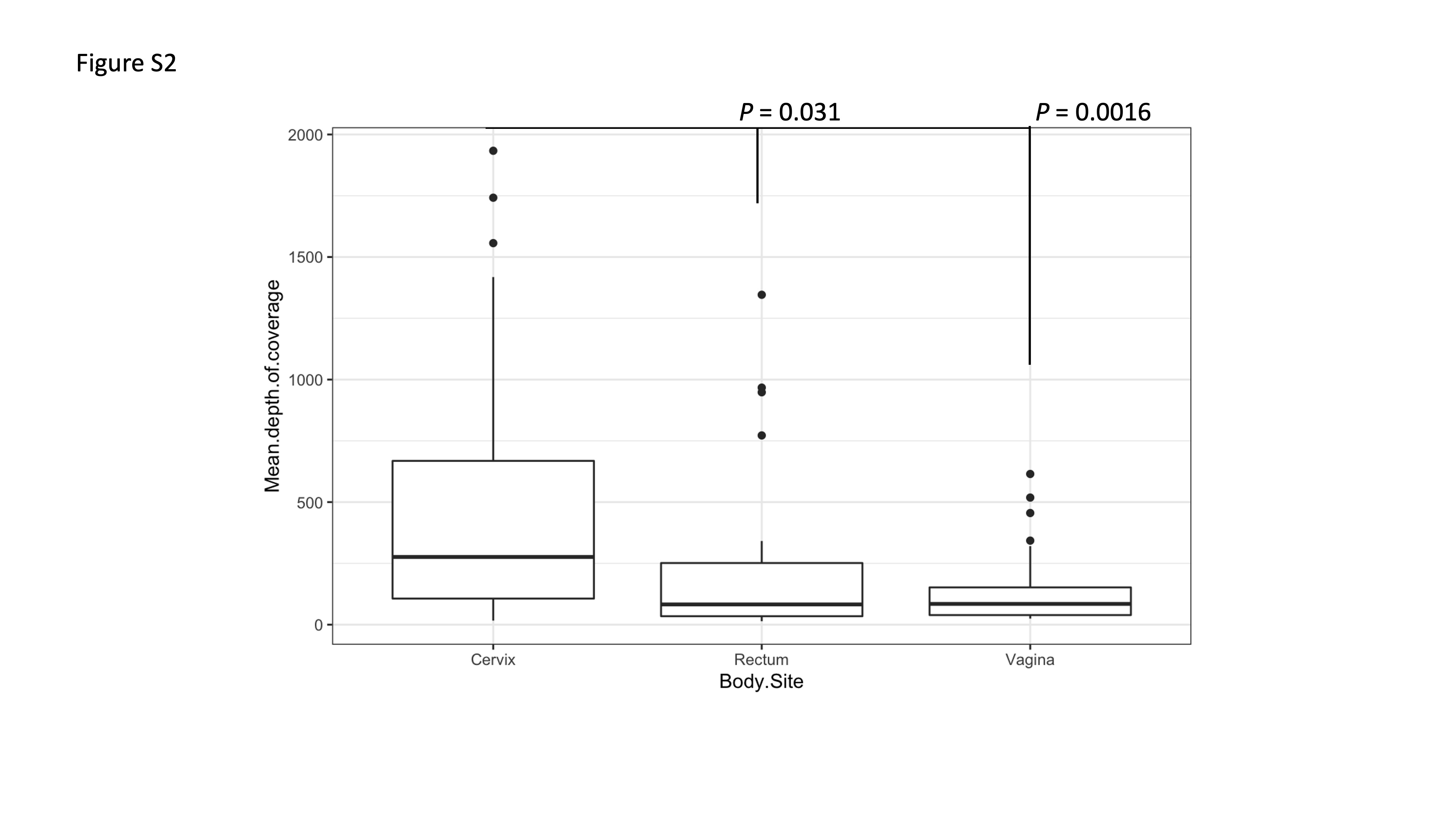

### Supplemental Figure 3

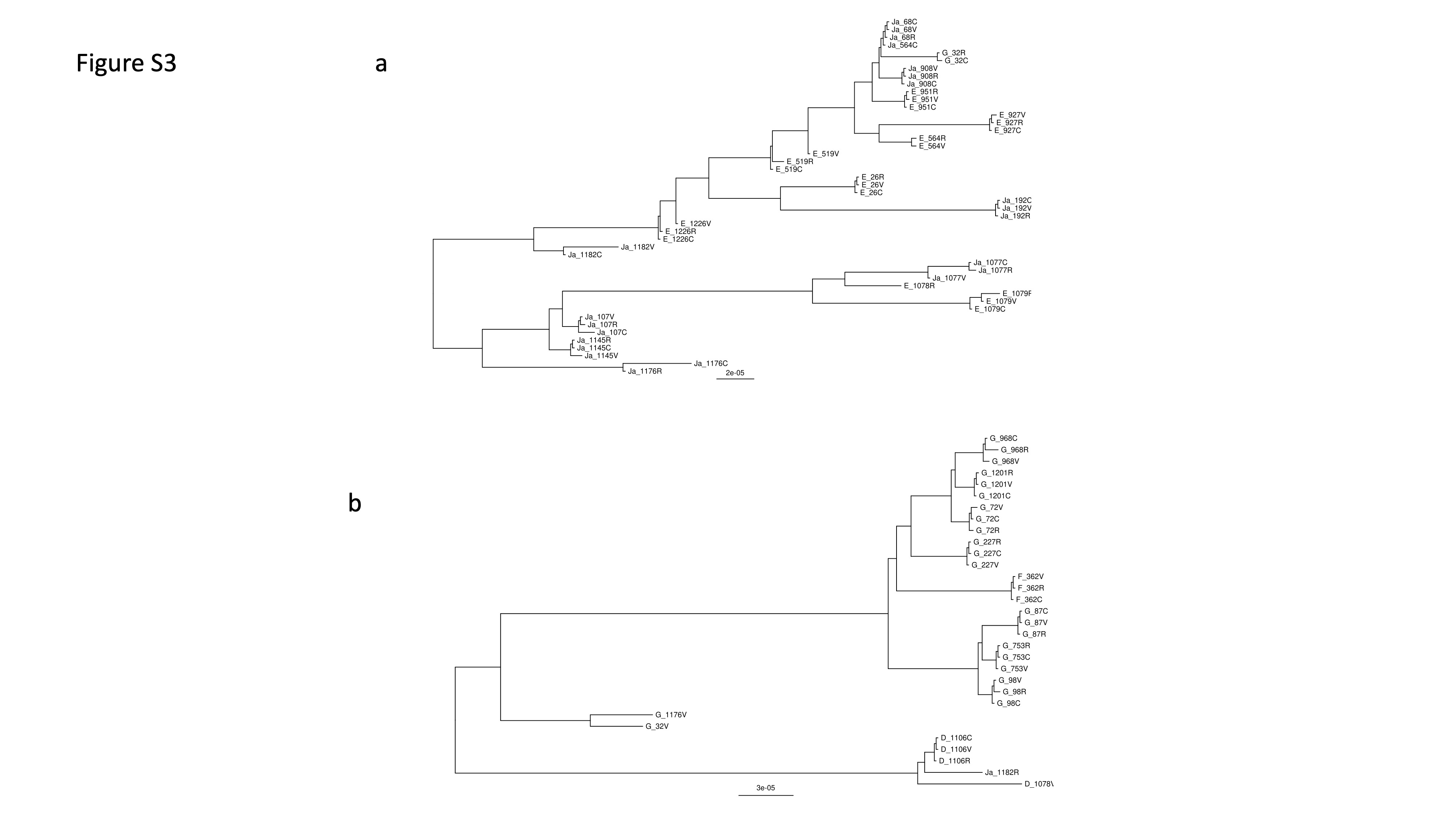

### Supplemental Figure 4

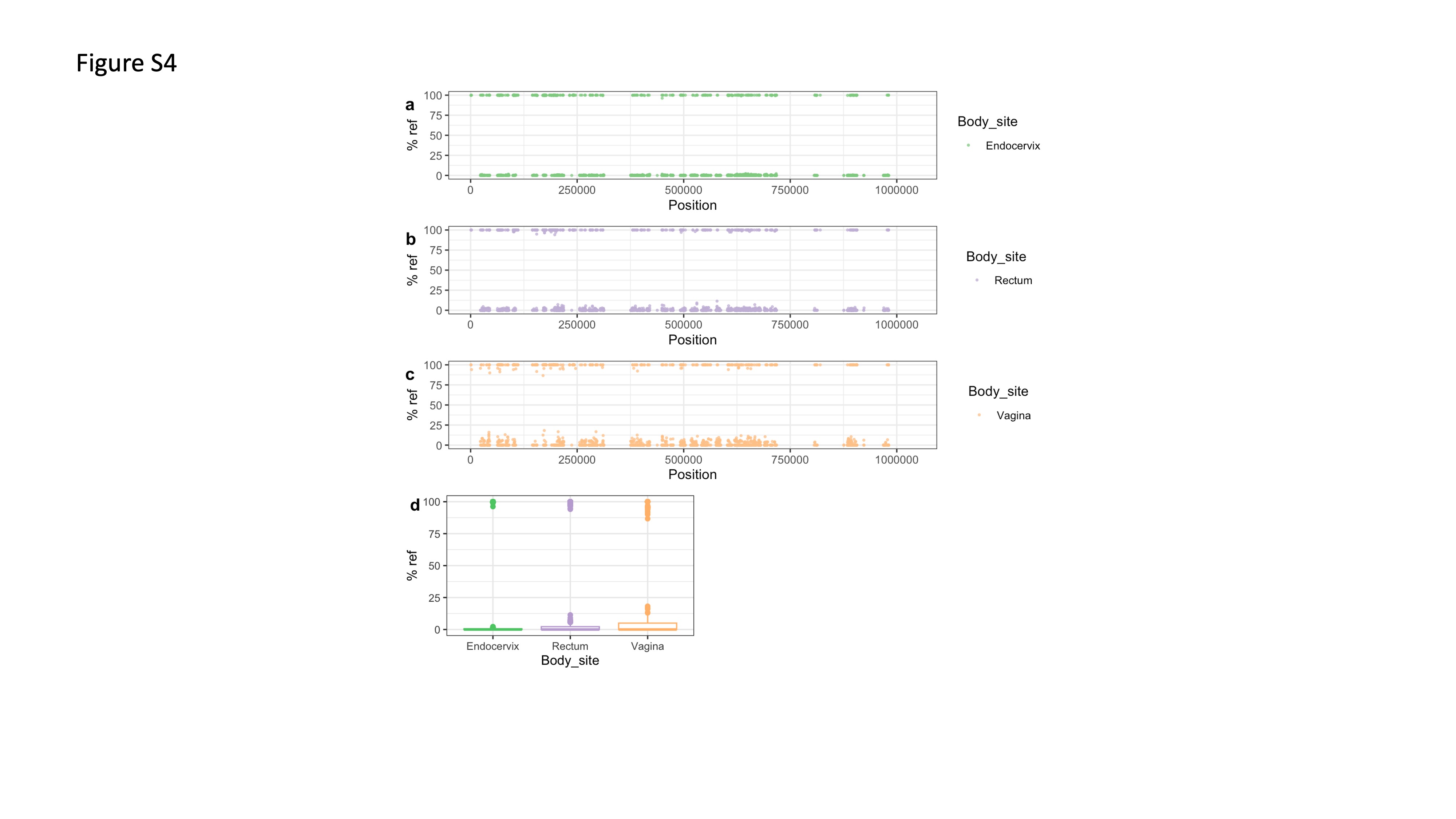

### Supplemental Figure 5

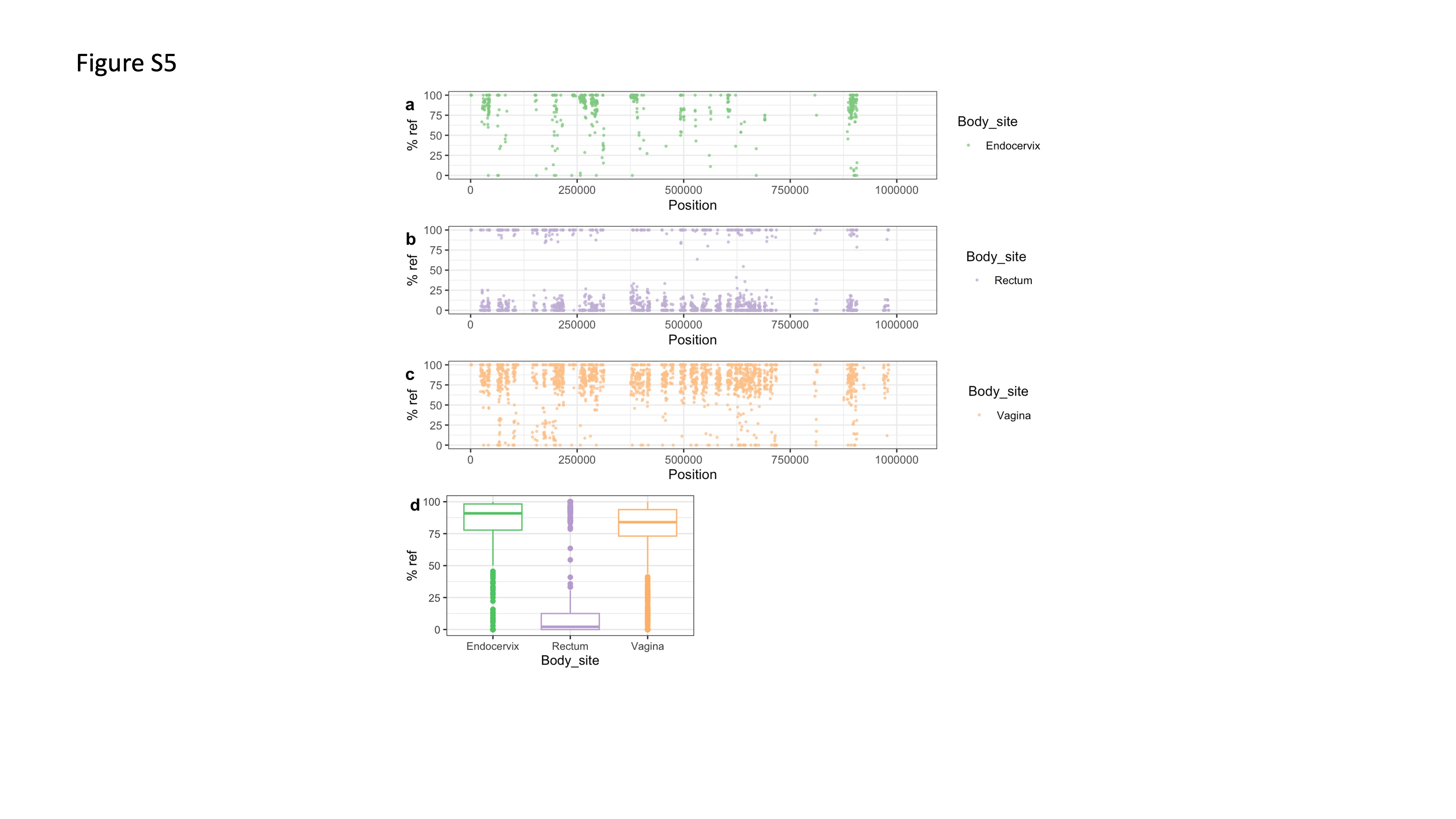
